# Cell strain energy costs of active control of contractility

**DOI:** 10.1101/2022.09.16.508225

**Authors:** Josephine Solowiej-Wedderburn, Carina M. Dunlop

## Abstract

Cell mechanosensing is implicated in the control of a broad range of cell behaviours, with cytoskeletal contractility a key component. Experimentally, it is observed that the contractility of the cell responds to increasing substrate stiffness, showing increased contractile force and changing the distribution of cytoskeletal elements. Here we show using a theoretical model of active cell contractility that upregulation of contractility need not be energetically expensive, especially when combined with changes in adhesion and contractile distribution. Indeed, we show that a feedback mechanism based on maintenance of strain energy would require an upregulation in contractile pressure on all but the softest substrates. We consider both the commonly reported substrate strain energy and active work done. We demonstrate substrate strain energy would select for the observed clustering of cell adhesions on stiffer substrates which also enable an upregulation of total contractile pressure; while localisation of contractility has the greatest impact on the internal work.

---

It is well established that cells sense, adapt and respond to the mechanical properties of their environment. This mechanosensing is key across cell behaviours ultimately affecting e.g. cell growth, development and differentiation [1–3]. Underpinning this mechanosensing are contractile cell forces generated by myosin motors within an actin rich network. These forces are transmitted from the cell to the extra cellular matrix (ECM) through cell adhesions [4]. A key focus of research into mechanotransduction has traditionally been these sites of cell adhesion, which in stiffer environments, are concentrated into small patches of strong attachment called focal adhesions [1, 3].

Experimental investigations into mechanosensing commonly use engineered gels or arrays of micropillars with defined mechanical properties to observe cell response to e.g. stiffer environments [5]. As well as changes in signal transduction, changes in the structural elements associated with cellular contractility are observed. These changes include increased actin density and stress fibre formation [6, 7]. Myosin has also been found to be more broadly distributed on soft gels becoming more localised overlapping with a dense actin cortex on stiffer substrates[8]. It is indeed typically observed that contractile forces increase in response to increased gel stiffness [9]. The mechanism by which stiffness change leads to a change in contractile force is unclear, although it has been hypothesised that contractility control could be based on a target stress or strain state [10].

Although changes in traction forces are typically measured in mechanostransduction studies, recent work suggests that substrate strain energy can be used as a measure for mechanical activity [11, 12]. It has been suggested that cells may respond to directly changes in substrate strain energy [13]. In [14] it was observed that substrate strain energy was approximately constant across a range of gel stiffness suggesting active control of strain energy. It is certainly clear that there are bounds on the energy budget of the cell [15], with a link between cell contractility and energy consumption demonstrated [16]. Theoretically models have explored this energy budget using energy constraints as drivers of differentiation [17] or cell shape control [18].

We here use an active matter model to investigate both the substrate strain energy and the work done by the active cell components, the active strain energy. We show that over a broad range of stiffnesses upregulation of active contractile pressure does not require an increase in energy expenditure. Indeed, upregulation of contractility is compatible with constant strain energy. We also investigate the localisation of contractility into the cortex of the cell showing that this will have minimal effect on the substrate strain energy at realistic levels of cell adhesion, although the active strain energy is sensitive to these changes. This is consistent with the observation that localisation of contractility can lead to large internal strains [19]. Introducing localised patches of adhesions we see that these generate polarised cells, with clustering of adhesion points particularly energetically favourable in terms of substrate strain energy, thus enabling significantly higher total contractile pressure to be generated.

## Model

Active matter models consider the cell as an elastic material with an additional component of stress as a result of active cellular contraction. Active contraction may be modelled either through computational simulations of cytoskeletal filaments [20–22] or via a continuum approach [23–25]. We adopt a continuum model taking the stress within the cell *σ* = *σ^P^* + *σ^A^*, where *σ^P^* is the passive cell elasticity and *σ^A^* is the active stress generated by the contractile pressure generated. Noting that the timescale for cell adhesion is faster than the relaxation timescale so that viscoelastic effects may be neglected we assume a linear elastic response in both the substrate and in the cell [25]. Furthermore, as the dimensions of the spread cell is greater than its height *h* we consider planar deformations only. The constitutive relation for stress is thus that 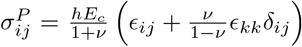, with *E_c_* the Young’s modulus of the cell and *ν* the Poisson’s ratio, and with 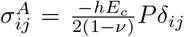, assuming isotropic contractility. The phenomenological function *P* couples the activity of the contractile machinery to the active stress. This function may be conceptualised in different ways. At a molecular level *P* may be expressed in terms of a chemical potential of ATP and its reaction products, which then drive the system out of equilibrium to generate active strains [26]. Alternatively we observe that when no forces are acting on the cell, including no cell adhesion, *P* = *ϵ_kk_*, which is a net area change so that *P* encompasses the idea of a ‘target’ strain of the cell system [10].

Cell contraction is resisted by adhesion to the underlying substrate and this force balance determines cell deformations. The in-plane force balance over the cell is

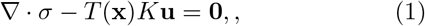

with zero stress imposed at the cell boundary. The resistance of the substrate to deformation is here modelled as –*K***u**, with *K* the substrate stiffness. This is a common first-order approximation for thin gel substrates [23, 27–29], where deformations are localized near the sites of applied traction generating an approximately linear relationship between stress and deformation. In the case of micropillar substrates [5] where the resisting stress is proportional to the pillar deflection the relationship is exact. To account for non-uniform adhesion of the cell to the substrate we set *T* (**x**) = 1 where the cell is adhered and *T* (**x**) = 0 where it is not. In the case of uniform adhesion (*T* (**x**) ≡ 1) we recover the force balance of [23, 27]. We here consider two further cases for *T* (**x**); a cell adhered over a ring and adhesions in spots mimicking cellular focal adhesion, imposing continuity of stress and deformation at internal boundaries.

There are two strain energies that we focus on. These are derived from the work done to the substrate *W_S_*; and the input of contractile work from network of myosin motors which (from the conversion of ATP) are assumed generate the active work done *W_CA_*. The substrate strain energy *W_S_* is easily experimentally measured and is often reported as a measure of mechanical activity, see e.g. [11–14]. The active work done *W_CA_* is also clearly important as it can be conceptually linked to the energy that is required to drive the cell into its contractile state, with hard constraints on its possible size [15]. However, *W_CA_* is significantly harder to directly experimentally quantify than *W_S_* and to interpret. Here, the energies *W_CA_* and *W_S_* can be calculated from the derived deformations as

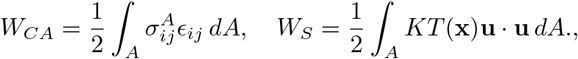

where the integrals are taken over the cell area *A*.

## Uniform and isotropic contractile pressure

We first consider that *P* = −*P*_0_ constant, so that the contractile pressure is uniform throughout the cell, as e.g. [23–30]. We consider two arrangements of adhesion: uniform cell adhesion to the substrate; and adhesion within a circular annulus. In these cases the cell deformations may be analytically determined, see Supporting Material for the full calculation, from which *W_S_* and *W_CA_* can also be analytically obtained.

In the case *T* (**x**) *≡* 1,

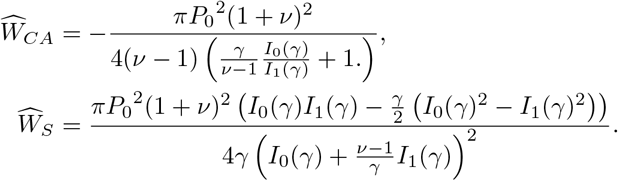

where 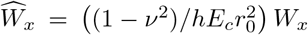, and *I_0,1_* are modified Bessel functions. 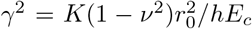 is the key nondimensional control parameter quantifying the relative stiffness of the cell and substrate for a cell of diameter 2*r*_0_, with stiffer gels corresponding to larger *γ*. These expressions relate the strain energies to the contractile pressure required to do the respective work.

Considering first the active strain energy *W_CA_*, from the result that *tI_N_* (*t*)/*I*_*N*+1_(*t*) is a monotone increasing function for *t* > 0 [31] we see as relative substrate stiffness increases (*γ* increases) *W_CA_* decreases in magnitude. Thus it is possible to upregulate the contractility strength *P*_0_ on stiffer substrates without increasing the energy expenditure. Indeed, if the energy *W_CA_* is constrained to be approximately constant, then this would necessitate an increase in contractility. In Fig. 1, we plot the contractility required to maintain 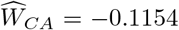 as substrate stiffness increases. The region above this curve corresponds to contractility that requires more energy, below this curve less energy is required despite the upregulation of contractility. Other energy contours demonstrate the same behaviour see Supplementary Figure S.1A.

**FIG. 1.**
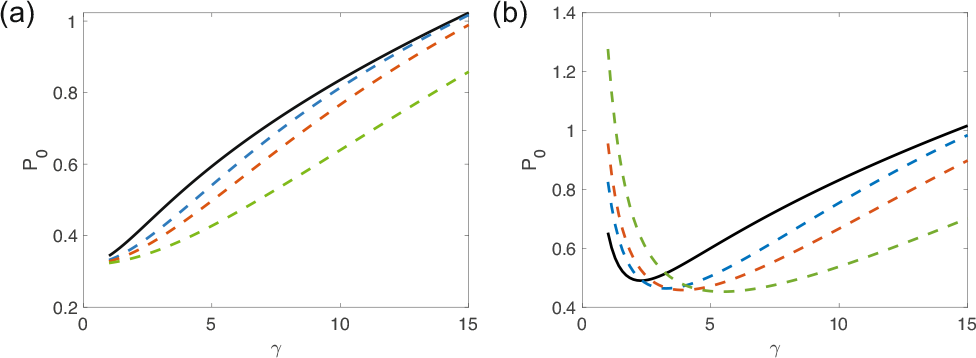
Lines of (a) constant 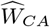 and (b) constant 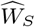 against increasing substrate stiffness (*γ* increasing), above each line more work is being done and below less. Solid black line is for complete adhesion and blue, orange, and green dashed lines are adhesion in outer rim at 30%, 20%, and 10% adhesion, respectively. (Corresponding to *r*_1_*/r*_0_ = 0.84, 0.89, 0.95, respectively). 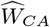 or 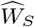 fixed values are, 0.1154 and 0.057, where we have chosen as the baseline the energy as for a completely adhered cell with *γ* = 7, *P*_0_ = 0.7.

For *W_S_* similar results hold for most values of *γ*. However, *W_S_* depends non-monotonically on *γ* with significantly different behaviour on the softest substrates. Looking at a fixed contour with 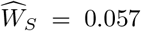, Fig. 1, then we see that, for example, for a completely adhered disc for values of *γ* > 2.3 increasing substrate stiffness enables greater contractile pressure with no extra work done. Note that again the region above the line corresponds to more work done and below to less work done than the baseline value. In contrast to *W_CA_*, we see that to increase, or even maintain, contractility on stiffer *γ* in the region of softest substrates (e.g. *γ* < 2.3, for a completely adhered disc) will require extra work. Other energy contours demonstrate the same behaviour see Supplementary Figure S.1B. This may be intuitively understood as there is need for greater contractility to generate the same work due to the inherently large deformations required on soft substrates.

Where cell adhesion is non-uniform but rather distributed in a ring around the cell edge so that *T* (*r*) = 1 in *r*_1_ < *r < r*_0_, similar analytical solutions may be obtained for *W_S_* and *W_CA_* in terms of *γ* and *P*_0_ (see Supporting Material). Similar qualitative behaviour is observed as for complete adhesion, see Fig. 1. Reducing the percentage of the cell adhered to the substrate reduces the amount of contractile pressure required to do the same work due to the lower resistance from the substrate on all but the softest substrates. However, the reduction in adhesion needs to be significant before this effect is seen with even an adhesive ring of width 20% of the cell radius still demonstrating very similar behaviour to complete adhesion, see Fig. 1. Notably, experimental results suggest that adhered area increases on stiffer substrates (e.g. [32] report adhered area increasing from 7% to 12% on 30kPa to 1MPa gels, see also [11, 33]). There is also a correlation between focal adhesion size and increased myosin activity e.g. [34]. Our model suggests that these results could be achieved without additional cost to the strain energy.

## Spatially varying contractile pressure

It is experimentally observed that the contractile apparatus of cells is concentrated towards the cell edge [35, 36]. Significantly the distribution of actin and myosin is observed to change in response to changes in stiffness, eventually concentrating in a thin more active cortex on stiff substrates [8]. To explore how altering the spatial distribution of contractile elements affects the cell energy budget we consider *P* = *P* (*r*), where *r* is a radial coordinate measured from the cell centre. We assume that *P* (*r*) is a monotonic decreasing function so that *P* generates greater contractile pressure at the cell edge. For specificity we take *P* (*r*) = − *a*(1 + *br^n^*), with *a, b* > 0 [19], which is chosen for analytical convenience, with *n* = 5 ensuring a strong differential in the contractile pressure. We take *a* = *P*_0_(*n* + 2)/(*n* + 2 + 2*br*_0_*^n^*) so that the total contractile pressure 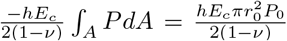 is the same as for uniform contractility. The parameter *b* adjusts the distribution of contractile elements, with increasing *b* corresponding to greater localisation of contraction to the cell edge. Provided the radial symmetry of the adhesion pattern is not broken an analytical solution is still available although more complex than that for uniform contractility, see Supporting Material.

In Fig. 2 (a)-(c), we set *P*_0_ = 0.7 (which is consistent with the contractile moment reported in [37]) and plot contours of fixed active energy as *γ* and *b* are varied. We see that increasing the substrate stiffness without altering the distribution of contractility or total contractile pressure will inevitably generate a reduction in *W_CA_* with any profile of contractility as expected. However we see by following contours that concentrating the contractility towards the edge of the cell (increasing *b*) can maintain *W_CA_*. This is observed across a range of adhesion percentages. Thus the concentration of contractility is playing a similar role to increased total contractility in terms of internal deformations and control of *W_CA_*. However, in considering the substrate strain energy *W_S_* a markedly different picture emerges, see Fig. 2 (d)-(f). Now depending on the localisation factor substrate strain energy may increase or decrease as substrate stiffness increases. In this way at lower values of the localisation factor the same nonlinear behaviour is observed as for uniform contractility, as might be expected. As localisation increases, we may need to either reduce or increase localisation to maintain a constant energy depending on the starting stiffness. Significantly, as the adhesion percentage reduces so the strain energy stored in the substrate becomes independent of the localisation of contractility with at 10% adhesion almost complete decoupling. Thus localisation as a mechanical component in mechanotransduction is likely most significant to internal stresses as has previously been suggested [19]. It is thus practically the case that the observed localisation of contractile machinery would not result in an increase in observed substrate strain energy, as is consistent with the observations of [14].

**FIG. 2.**
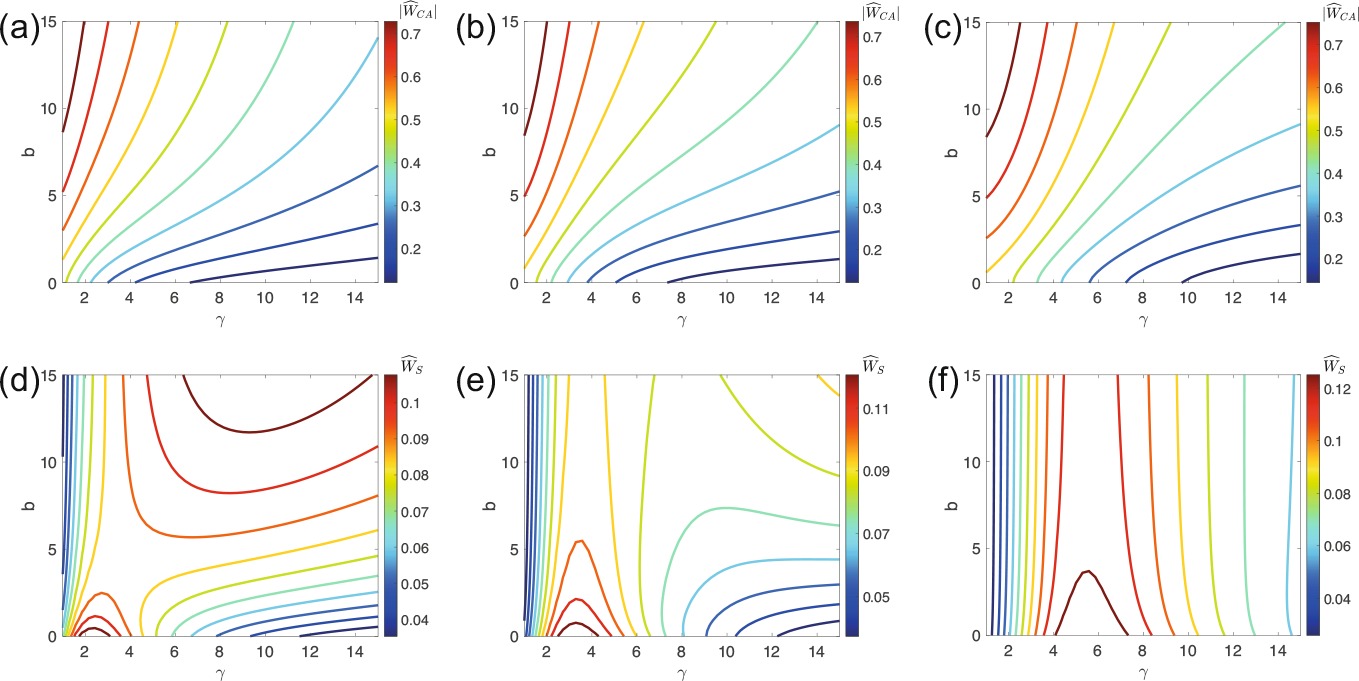
(a)–(c) Contour plots of fixed 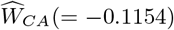: (a) complete adhesion; (b) adhered ring covering 30% adhered area; (c) adhered ring covering 10% adhered area. (d)–(f) Contour plots of fixed 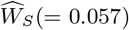: (d) complete adhesion; (e) adhered ring covering 30% adhered area; (f) adhered ring covering 10% adhered area. *P*_0_ = 0.7 for all plots.

## Adhesion patterning

We now consider adhesion in small localised patches to investigate the role of focal adhesions in the work done by the cell. We consider a pattern of 20 spots either evenly distributed around the cell edge (Fig. 3(a)); or alternatively in two clusters at opposite ends of the cell to generate polarised cell shapes (Fig. 3(b)). In the case of distinct adhered spots, analytical solutions cannot be obtained and we solve Eq.(1) finite element methods implemented in MATLAB 2019b (PDE Toolbox), see Supplementary Material for numerical details.

**FIG. 3.**
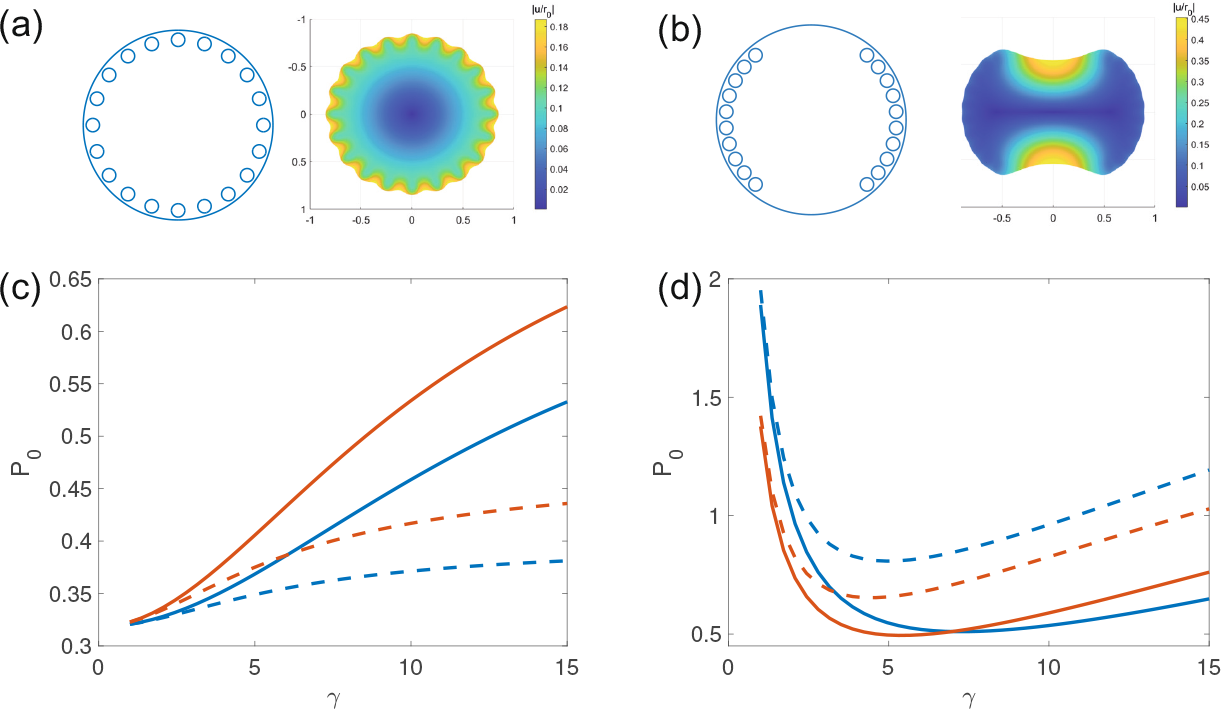
(a) An adhesion pattern with 20 spots evenly distributed around the cell edge (10% adhered area) and the corresponding cell deformation (*γ* = 7 and *P*_0_ = 0.7). (b) An adhesion pattern with 20 spots distributed in two clusters and the resultant deformation (*γ* = 7 and *P*_0_ = 0.7). (c) and (d) Plots of total cellular contractility *P*0 that maintain (c) fixed *W_CA_* 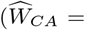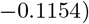 and (d) fixed 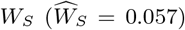 for adhesion patterns of 20 spots distributed either evenly (solid lines) or in two clusters (dashed lines) for two different spot sizes - 5% overall adhered area in blue, and 10% overall adhered area in orange.

As in the case of an adhered ring, for a given spot distribution *W_CA_* would naturally reduce on stiffer substrates while *W_S_* has the same initial increase over the softest values and then reduction on stiffer, Fig. 3(c) and (d). As such, we see that over much of the stiffness range contractility can increase without energy penalty and indeed any feedback mechanism based on constraining energy changes would necessitate an increase in contractility. We observe significant differences in the effects of spot clustering on *W_CA_* and *W_S_*. Clustering of adhesions increases *W_CA_* as larger cell deformations are incurred so that the possible contractility before energy increase is reduced (dashed line) reducing the possible increases in contractility. However, adhesion clustering reduces *W_S_* so that this arrangement is energetically favourable in terms of substrate energy, as reported previously [30].

Significantly, experimental results suggest that adhesion distribution changes on stiffer substrates, resulting in more polarised cell shapes [37, 38]. This experimentally observed behaviour would be energetically favourable under a consideration of *W_S_*, Fig. 3 (d), enabling greater upregulation of contractility, particularly on stiffer substrates. This further supports the substrate strain energy as a useful correlative measure of mechanical activity, as suggested by e.g. [13]. Additionally, we see that if we constrain substrate strain energy as has been reported [14], this requires an upregulation of contractile activity on stiffer substrates.

In conclusion, we have demonstrated both analytically and numerically that upregulating contractility on stiffer substrates need not be energetically expensive. Indeed under constraints of maintaining cell energy it is necessary for contractile pressure to increase to prevent a reduction in either the active energy or substrate energy on stiffer substrates. We note that the observed localisation of contractility to the cell edge has only limited influence on substrate strain energy at realistic adhesion percentages, although such localisation does change the internal work done, in part due to the large internal strains that can be generated [19]. Significantly, we show that clustering of cell adhesions on stiffer substrates generate polarised cell shapes which are energetically favourable in terms of substrate strain energy, and which enable significant contractile upregulation.

## ACKNOWLEDGMENTS

J.S.-W. acknowledges PhD funding from the UK EPSRC, Institutional Doctoral Training Partnership (EP/N509772/1) and postdoctoral EPSRC funding (EP/W522302/1). C.M.D. also acknowledges financial support from the UK EPSRC (EP/M012964/1).

