## Supplementary Data for "Cell strain energy costs of active control of contractility"

### Supplementary Material: Cell strain energy costs of active control of contractility

J. Solowiej-Wedderburn and C. M. Dunlop

#### Figure S1

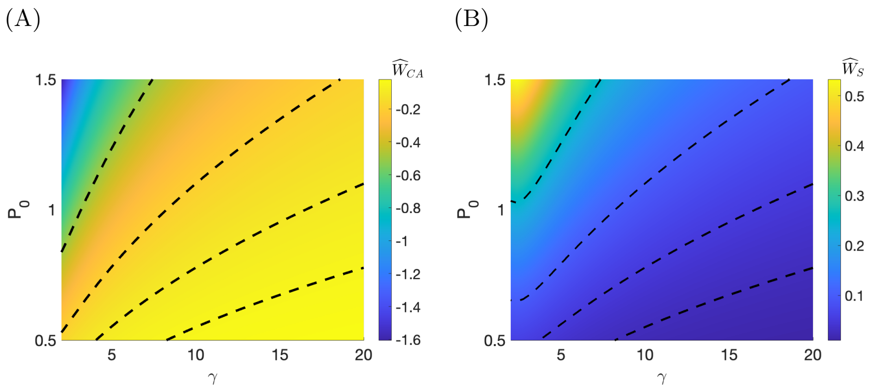


Strain energies for a completely adhered cell with uniform contractile pressure $P_0$ on substrate of stiffness $\gamma$: (A) Active strain energy $\widehat{W}_{CA}$, black dashed lines contours at $\widehat{W}_{CA} = -0.5, -0.2, -0.1, -0.05$ from top to bottom. (B) Substrate strain energy $\widehat{W}_S$, black dashed lines contours at $\widehat{W}_S = 0.25, 0.1, 0.05, 0.025$ from top to bottom.

#### S1 Derivation of analytical solutions of the main force balance equations

The force balance equation that is solved to determine the cell deformations is

$$\nabla \cdot \sigma - KT(\mathbf{x})\mathbf{u} = \mathbf{0}, \tag{1}$$

where $T(\mathbf{x}) = 1$ where the cell is adhered and $T(\mathbf{x}) = 0$ where there is no adhesion. We impose zero stress on the cell outer boundary and continuity of stress and deformation at all internal boundaries. The parameter $K$ as defined in the main paper quantifies the stiffness of the substrate. The constitutive relationship relating stress and strain is taken as

$$\sigma_{ij} = \frac{hE_c}{1+\nu}\left(\epsilon_{ij} + \frac{\nu}{1-\nu}\epsilon_{kk}\delta_{ij}\right) - \frac{hE_c}{2(1-\nu)}P\delta_{ij}, \tag{2}$$

where we have assumed plane stress, linear elasticticy and isotropic contractile pressure. The parameters are $E_c$, the Young's modulus of the cell, $\nu$, the Poisson's ratio and $h$ the height of the cell, which we assume is initially circular with radius $r_0$. The function $P$ models the contractility of the cell and we consider both a constant pressure and a contractile pressure that is monotonically increasing in magnitude.

We here derive the analytical solutions for the deformations and strain energies presented in the paper under the various assumptions on adhesion and contractile pressure.

##### Case 1: Uniform contractile pressure $P = -P_0$ and complete adhesion of cell to substrate $T(\mathbf{x}) \equiv 1$

We first consider the paradigm case of a cell completely adhered across the cell-substrate interface with a uniform isotropic contractile pressure $P = -P_0$. In this case, $T(\mathbf{x}) \equiv 1$. Using the radial symmetry of the problem to note that the deformations are purely radial such that $\mathbf{u} = u(r)\mathbf{e_r}$ in Eq. (2), Eq. (1) then becomes

$$\hat{r}^2 \frac{d^2 \hat{u}}{d\hat{r}^2} + \hat{r}\frac{d\hat{u}}{d\hat{r}} - \left(1 + \gamma^2 \hat{r}^2\right) \hat{u} = 0, \tag{3}$$

where length scales have been normalised by the cell radius ($\hat{r} = r/r_0$ and $\hat{u} = u/r_0$) and $\gamma$ is a dimensionless parameter such that $\gamma^2 = K(1-\nu^2)r_0^2/hE_c$, see also e.g. [1, 2].

We solve equation (3) with boundary conditions $\hat{u}(\hat{r} = 0) = 0$ and no stress $\sigma \cdot \mathbf{n} = \mathbf{0}$ at the outer edge $\hat{r} = 1$. With $P = -P_0$ this no stress condition becomes (using Eq. 2))

$$\left.\frac{d\hat{u}}{d\hat{r}}\right|_{\hat{r}=1} + \nu\hat{u}(\hat{r} = 1) = \frac{-P_0(1+\nu)}{2}.$$

Equation (3) is the modified Bessel Equation [3] and so we find the cell deformation in terms of modified Bessel functions as

$$\hat{u} = \frac{-P_0(1+\nu)}{2\left(\gamma I_0(\gamma) - (1-\nu)I_1(\gamma)\right)} I_1(\gamma\hat{r}), \tag{4}$$

as in [1].

The active strain energy is defined as as

$$W_{CA} = \frac{1}{2}\int_A \sigma_{ij}^A \epsilon_{ij}\, dA,$$

and the substrate strain energy as

$$W_S = \frac{1}{2}\int_A KT(\mathbf{x})\mathbf{u}\cdot\mathbf{u}\, dA.$$

Consistent with our rescaling of the force balance equation (1), we introduce the dimensionless stress $\widehat{\sigma}_{ij} = \left((1-\nu^2)/hE_c\right)\sigma_{ij}$ and rescale the energies as $\widehat{W}_x = \left((1-\nu^2)/hE_c r_0^2\right) W_x$. As such, we have the dimensionless energy expressions

$$\widehat{W}_{CA} = \frac{1}{2}\int_{\widehat{A}} \widehat{\sigma}_{ij}^A \epsilon_{ij}\, d\widehat{A}, \tag{5}$$

$$\widehat{W}_S = \frac{1}{2}\int_{\widehat{A}} \gamma^2 T(\hat{\mathbf{x}})\hat{\mathbf{u}}\cdot\hat{\mathbf{u}}\, d\widehat{A}, \tag{6}$$

where $\widehat{A}$ is the normalised cell area $\pi$.

Substituting the expression (4) into (5) and (6) and integrating we determine expressions for the active strain energy and work done to the substrate by a completely adhered disc with uniform isotropic contractility.

$$
\begin{aligned}
\widehat{W}_{CA} &= \frac{\pi P_0(1+\nu)}{2}\int_0^1 \varepsilon_{ii} r dr \\
&= -\frac{\pi P_0^2(1+\nu)^2}{4\gamma\left(\gamma I_0(\gamma)-(1-\nu)I_1(\gamma)\right)}\int_0^\gamma I_0(z) z dz \\
&= -\frac{\pi P_0^2(1+\nu)^2}{4(\nu-1)\left(\frac{\gamma}{(\nu-1)}\frac{I_0(\gamma)}{I_1(\gamma)}+1\right)},
\end{aligned}
\tag{7}
$$

$$
\begin{aligned}
\widehat{W}_{S} &= \frac{\pi P_0^2(1+\nu)^2}{4\left(\gamma I_0(\gamma)-(1-\nu)I_1(\gamma)\right)^2}\int_0^\gamma I_1(z)^2 z dz \\
&= \frac{\pi P_0^2(1+\nu)^2\left(I_0(\gamma)I_1(\gamma)-\frac{\gamma}{2}\left((I_0(\gamma))^2-(I_1(\gamma))^2\right)\right)}{4\gamma\left(I_0(\gamma)+\frac{(\nu-1)}{\gamma}I_1(\gamma)\right)^2},
\end{aligned}
\tag{8}
$$

where we have used integration by parts to find

$$
\begin{aligned}
\int_0^\gamma I_1(z)^2 z dz &= \left[z I_0(z) I_1(z)\right]_0^\gamma - \int_0^\gamma I_0(z)^2 z dz, \\
\int_0^\gamma I_0(z)^2 z dz &= \left[\frac{1}{2}z^2 I_0(z)^2\right]_0^\gamma - \int_0^\gamma z^2 I_0(z) I_1(z) dz, \\
\int_0^\gamma z^2 I_0(z) I_1(z) dz &= \left[\frac{1}{2}z^2 I_1(z)^2\right]_0^\gamma,
\end{aligned}
$$

see also [3] for the integrals of modified Bessel functions.

#### Case 2: Uniform contractile pressure $P = -P_0$ and adhesion of cell to substrate in an outer annulus

We now consider a cell with uniform isotropic contractile pressure, but adhesion restricted to an annular region around the cell edge. Now $T(r) = 1$ on $r_1 < r < r_0$. As previously, we normalise length scales by the cell radius such that $T(\widehat{r}) = 1$ on $\widehat{r}_1 < \widehat{r} < 1$ with $\widehat{r}_1 = r_1/r_0$. The problem is still radially symmetric. We substitute the adhesion geometry into the force balance equation (1). We apply the same scaling as in the paradigm case dropping the hats here and all in further sections. Hence, (1) becomes

$$
r^2\frac{d^2u}{dr^2} + r\frac{du}{dr} - u = 0, \quad \text{on } 0 < r < r_1 \tag{9}
$$

$$
\& \quad r^2\frac{d^2u}{dr^2} + r\frac{du}{dr} - \left(1+\gamma^2 r^2\right)u = 0, \quad \text{on } r_1 \le r < 1, \tag{10}
$$

with a no stress boundary condition at $r = 1$; $u(r = 0) = 0$; and continuity in both cell deformation and stress across the internal boundary at $r = r_1$. We solve (9)–(10) to find

$$u = \begin{cases} P_0\, \alpha_0\, r, & r \in [0, r_1) \\ P_0\left(\alpha_1 I_1(\gamma r) + \beta_1 K_1(\gamma r)\right), & r \in [r_1, 1], \end{cases} \tag{11}$$

where $I_1$ and $K_1$ are modified Bessel functions, as in [2]. The constants $\alpha_0 = \frac{1}{r_1}\left(\alpha_1 I_1\left(\gamma r_1\right) + \beta_1 K_1\left(\gamma r_1\right)\right)$, $\alpha_1 = -\frac{(1+\nu)}{2\gamma}\cdot\left(F(\gamma) - G(\gamma)H\left(\gamma r_1\right)\right)^{-1}$, $\beta_1 = H\left(\gamma r_1\right)\alpha_1$, and, for simplicity, we have introduced

$$\begin{aligned} F(z) &= I_0(z) + \frac{(\nu - 1)}{z} I_1(z), \\ G(z) &= K_0(z) - \frac{(\nu - 1)}{z} K_1(z), \\ H(z) &= \left(\frac{z I_0(z) - 2 I_1(z)}{z K_0(z) + 2 K_1(z)}\right). \end{aligned}$$

We substitute expression (11) for the cell deformation into (5) to determine an analytic expression for the active strain energy of a cell with an adhered ring as

$$\begin{aligned} W_{CA} &= \frac{(1+\nu)P_0}{4} \int_0^{2\pi} \int_0^1 \varepsilon_{ii} r\, dr\, d\theta \\ &= \frac{\pi P_0^2 (1+\nu)}{2} \Big( \alpha_0 r_1^2 + \alpha_1 \left[ I_1(\gamma) - r_1 I_1\left(\gamma r_1\right) \right] + \beta_1 \left[ K_1(\gamma) - r_1 K_1\left(\gamma r_1\right) \right] \Big). \end{aligned} \tag{12}$$

Similarly, substituting (11) into (6) we determine an analytic expression for the work done to the substrate by a cell with an adhered ring as

$$
\begin{aligned}
W_S &= \frac{\gamma^2}{2}\int_0^{2\pi}\int_{r_1}^{1} u^2 r dr d\theta \\
&= P_0^2\gamma^2\pi\int_{r_1}^{1}\left(\alpha_1 I_1(\gamma r)+\beta_1 K_1(\gamma r)\right)^2 r dr \\
&= P_0^2\pi\Bigg(\alpha_1^2\bigg[\gamma I_0(\gamma)I_1(\gamma)-\frac{\gamma^2}{2}\left(I_0(\gamma)^2-I_1(\gamma)^2\right) \\
&\qquad\qquad -\gamma r_1 I_0\left(\gamma r_1\right)I_1\left(\gamma r_1\right)+\frac{\gamma^2 r_1^2}{2}\Big(I_0\left(\gamma r_1\right)^2-I_1\left(\gamma r_1\right)^2\Big)\bigg] \\
&\qquad +2\alpha_1\beta_1\bigg[\gamma I_0(\gamma)K_1(\gamma)+\frac{\gamma^2}{2}\left[I_0(\gamma)K_0(\gamma)+I_1(\gamma)K_1(\gamma)\right] \\
&\qquad\qquad -\gamma r_1 I_0\left(\gamma r_1\right)K_1\left(\gamma r_1\right)-\frac{\gamma^2 r_1^2}{2}\Big(I_0\left(\gamma r_1\right)K_0\left(\gamma r_1\right)+I_1\left(\gamma r_1\right)K_1\left(\gamma r_1\right)\Big)\bigg] \\
&\qquad -\beta_1^2\bigg[\gamma K_0(\gamma)K_1(\gamma)+\frac{\gamma^2}{2}\left(K_0(\gamma)^2-K_1(\gamma)^2\right) \\
&\qquad\qquad -\gamma r_1 K_0\left(\gamma r_1\right)K_1\left(\gamma r_1\right)-\frac{\gamma^2 r_1^2}{2}\Big(K_0\left(\gamma r_1\right)^2-K_1\left(\gamma r_1\right)^2\Big)\bigg]\Bigg),
\end{aligned}
\tag{13}
$$

where we have evaluated the integral expressions

$$
\begin{aligned}
\int zI_1(z)^2dz &= zI_0(z)I_1(z)-\frac{z^2}{2}\left(I_0(z)^2-I_1(z)^2\right), \\
\int zK_1(z)^2dz &= -zK_0(z)K_1(z)-\frac{z^2}{2}\left(K_0(z)^2-K_1(z)^2\right), \\
\int zI_1(z)K_1(z)dz &= zI_0(z)K_1(z)+\frac{z^2}{2}\left(I_0(z)K_0(z)+I_1(z)K_1(z)\right)
\end{aligned}
$$

using integration by parts.

As in the case of complete adhesion, we see that with a uniform isotropic contractility the expressions for $\widehat{W}_{CA}$ and $\widehat{W}_S$ are proportional to $P_0^2$ and so the contractility required to maintain each energy constant may be determined by rearranging (12) and (13).

#### Case 3: Strain energies for a completely adhered cell with differential contractility

For a completely adhered circular cell with contractility given by $P(r) = -a(1+br^5)$, the force balance equation (1) becomes (where again we are using the scaled nondimensional variables but have dropped the hats)

$$r^2\frac{d^2u}{dr^2} + r\frac{du}{dr} - (1+\gamma^2r^2)u = -\frac{5ab}{2}(1+\nu)r^6, \tag{14}$$

with boundary conditions

$$u(r=0) = 0, \tag{15}$$

$$\left.\frac{du}{dr}\right|_{r=1} + \nu u(r=1) = -\frac{(1+\nu)a(1+b)}{2}. \tag{16}$$

We solve the system given by (14)–(16) using the method of variation of parameters to find

$$u(r) = \frac{(1+\nu)}{2\gamma}\left(cI_1(\gamma r) + \frac{5ab}{\gamma^5}\left[\gamma^4r^4 + 15\gamma^2r^2 - \frac{45\pi}{2}L_1(\gamma r)\right]\right), \tag{17}$$

where we have defined

$$\begin{aligned} c := \frac{-a(1+b)}{F(\gamma)} + \frac{5ab}{\gamma^5F(\gamma)}\Bigg[&(1-\nu)\left(\gamma^3 + 15\gamma - \frac{45\pi}{2\gamma}L_1(\gamma)\right) \\ &- \left(5\gamma^3 + 45\gamma - \frac{45\pi}{2}L_0(\gamma)\right)\Bigg], \end{aligned} \tag{18}$$

given in terms of the modified Struve functions $L_0(z)$ and $L_1(z)$ [3]. Recall, function $F(\gamma)$ was defined by

$$F(z) := I_0(z) + \frac{(\nu-1)}{z}I_1(z).$$

We may substitute the expression for cell deformation (17) into (5) to determine the active strain energy

$$
\begin{aligned}
W_{CA} &= \frac{1}{2}\int_A \sigma_{ij}^A \varepsilon_{ij} dA \\
&= \frac{(1+\nu)}{2}\pi a \int_0^1 \left(1+br^5\right)\left(\frac{du}{dr}+\frac{u}{r}\right) r dr \\
&= \frac{(1+\nu)^2}{4\gamma}\pi a \int_0^1 \left(1+br^5\right)\left(c\gamma r I_0(\gamma r) + \frac{5ab}{\gamma^5}\left[5\gamma^4 r^4 + 45\gamma^2 r^2 - \frac{45\pi}{2}\gamma r L_0(\gamma r)\right]\right) dr \\
&= \frac{(1+\nu)^2}{4\gamma}\pi a \Bigg( c\gamma \left[\int_0^1 r I_0(\gamma r) dr + b\int_0^1 r^6 I_0(\gamma r) dr\right] \\
&\qquad + \frac{5ab}{\gamma^5}\Bigg[5\gamma^4 \int_0^1 r^4 dr + 5b\gamma^4 \int_0^1 r^9 dr + 45\gamma^2 \int_0^1 r^2 dr + 45b\gamma^2 \int_0^1 r^7 dr \\
&\qquad - \frac{45\pi}{2}\int_0^1 \gamma r L_0(\gamma r) dr - \frac{45\pi b\gamma}{2}\int_0^1 \gamma r^6 L_0(\gamma r) dr \Bigg) \\
&= \frac{(1+\nu)^2}{4\gamma}\pi a \Bigg( \frac{5ab}{\gamma^5}\left[\gamma^4 + 15\gamma^2 + b\left(\frac{\gamma^4}{2}+\frac{45\gamma^2}{8}\right)\right] + cI_1(\gamma) - \frac{225\pi ab}{2\gamma^5} L_1(\gamma) \\
&\qquad + cb\gamma \int_0^1 r^6 I_0(\gamma r) dr - \frac{225\pi ab^2}{2\gamma^4}\int_0^1 r^6 L_0(\gamma r) dr \Bigg). \qquad (19)
\end{aligned}
$$

We explicitly evaluate the integral expressions

$$
\begin{aligned}
\int_0^1 r^6 I_0(\gamma r) dr &= \frac{1}{\gamma^6}\Bigg[\left(\gamma^5 + 25\gamma^3 + 225\gamma - \frac{225\pi}{2}L_0(\gamma)\right) I_1(\gamma) \\
&\qquad - \left(5\gamma^4 + 75\gamma^2 - \frac{225\pi}{2}L_1(\gamma)\right) I_0(\gamma)\Bigg], \\
\int_0^1 r^6 L_0(\gamma r) dr &= \frac{1}{\gamma^5}\Bigg[\gamma^4 L_1(\gamma) + 25\gamma^2 L_1(\gamma) + 225 L_1(\gamma) - 5\gamma^3 L_0(\gamma) - 75\gamma L_0(\gamma) \\
&\qquad - \frac{225}{\pi}\, {}_2F_3\left(1,1;\frac{1}{2},\frac{3}{2},2;\frac{\gamma^2}{4}\right) + \frac{5\gamma^4}{3\pi} + \frac{75\gamma^2}{2\pi} + \frac{225}{\pi}\Bigg],
\end{aligned}
$$

where ${}_pF_q(a_1,...,a_p;b_1,...,b_q;z)$ is the generalised hypergeometric function [3, 4].

Similarly, substituting (17) into (6) we determine the work done to the substrate by a completely adhered cell with differential contractility as

$$
\begin{aligned}
W_S &= \frac{1}{2}\int_A \gamma^2 \mathbf{u}\cdot\mathbf{u}dA \\
&= \gamma^2\pi\int_0^1 u^2 r dr \\
&= \frac{\pi(1+\nu)^2}{4\gamma^2}\int_0^\gamma \left[c^2 z I_1(z)^2 + \frac{10abc}{\gamma^5}\left(z^5 I_1(z) + 15z^3 I_1(z) - \frac{45\pi}{2} z I_1(z) L_1(z)\right)\right. \\
&\qquad + \frac{25a^2b^2}{\gamma^{10}}\left(z^9 + 30z^7 + 225z^5 - 45\pi z^5 L_1(z)\right. \\
&\qquad\qquad \left.\left. - 675\pi z^3 L_1(z) + \left(\frac{45\pi}{2}\right)^2 z L_1(z)^2\right)\right] dz,
\end{aligned}
$$

which we re-write as

$$
W_S = c^2 N_1(\gamma) + abcN_2(\gamma) + a^2b^2 N_3(\gamma), \tag{20}
$$

where we have defined

$$
\begin{aligned}
N_1(\gamma) &:= \frac{\pi(1+\nu)^2}{4\gamma^2}\int_0^\gamma z I_1(z)^2 dz = \frac{\pi(1+\nu)^2}{4\gamma}\left(I_0(\gamma)I_1(\gamma) - \frac{1}{2}\gamma\left(I_0(\gamma)^2 - I_1(\gamma)^2\right)\right), \\
N_2(\gamma) &:= \frac{5\pi(1+\nu)^2}{2\gamma^7}\int_0^\gamma \left(z^5 I_1(z) + 15z^3 I_1(z) - \frac{45\pi}{2} z I_1(z) L_1(z)\right) dz \\
&= \frac{5\pi(1+\nu)^2}{2\gamma^6}\left[\left(\gamma^4 + 30\gamma^2 - \frac{315\pi}{4}L_1(\gamma) + \frac{45\pi}{4}\gamma L_0(\gamma)\right) I_0(\gamma)\right. \\
&\qquad \left. - \left(5\gamma^3 + \frac{225}{2}\gamma - \frac{225\pi}{4}L_0(\gamma) + \frac{45\pi}{4}\gamma L_1(\gamma)\right) I_1(\gamma)\right], \\
N_3(\gamma) &:= \frac{25\pi(1+\nu)^2}{4\gamma^{12}}\left(\frac{1}{10}\gamma^{10} + \frac{15}{4}\gamma^8 + \frac{75}{2}\gamma^6\right. \\
&\qquad \left. + \int_0^\gamma \left(\left(\frac{45\pi}{2}\right)^2 z L_1(z)^2 - 45\pi z^5 L_1(z) - 675\pi z^3 L_1(z)\right) dz\right.,
\end{aligned}
$$

with

$$\int_0^\gamma zL_1(z)^2dz = \gamma L_0(\gamma)L_1(\gamma) + \frac{1}{2}\gamma^2\left(L_1(\gamma)^2 - L_0(\gamma)^2\right)$$
$$+\frac{2}{\pi}\left(\gamma^2L_1(\gamma) - 2\gamma L_0(\gamma) + \frac{2\gamma^2}{\pi} + \frac{2\gamma^2}{\pi}\,{}_2F_3\left(1,1;\frac{3}{2},\frac{3}{2},2;\frac{\gamma^2}{4}\right)\right),$$
$$\int_0^\gamma z^5L_1(z)dz = -\frac{1}{\pi}\left(\frac{1}{3}\gamma^6 + \frac{15}{2}\gamma^4 + 45\gamma^2\right) + \left(\gamma^5 + 15\gamma^3 + 45\gamma\right)L_0(\gamma)$$
$$-5\left(\gamma^4 + 9\gamma^2\right)L_1(\gamma) - \frac{45\gamma^2}{\pi}\,{}_2F_3\left(1,1;\frac{3}{2},\frac{3}{2},2;\frac{\gamma^2}{4}\right),$$
$$\int_0^\gamma z^3L_1(z)dz = -\frac{1}{2\pi}\left(\gamma^4 + 6\gamma^2\right) + \left(\gamma^3 + 3\gamma\right)L_0(\gamma) - 3\gamma^2L_1(\gamma) - \frac{3\gamma^2}{\pi}\,{}_2F_3\left(1,1;\frac{3}{2},\frac{3}{2},2;\frac{\gamma^2}{4}\right).$$

We can see from expression (18) that $c$ is directly proportional to $a$ and of the form $c = xb + y$. Hence, we see that the expressions for $W_{CA}$ and $W_S$ are proportional to $a^2$ and quadratic in $b$ which may be used to determine the differential contractility profile required to maintain a particular constraint on either $W_{CA}$ or $W_S$.

#### Strain energies for an adhered annulus with differential contractility

For a circular cell with and adhered ring and differential contractility given by $P(r) = -a(1 + br^5)$, the force balance equation (1) becomes (again we are using the scaled nondimensional variables but have dropped the hats)

$$r^2\frac{d^2u}{dr^2} + r\frac{du}{dr} - u = -\frac{5ab}{2}(1+\nu)r^6, \quad \text{on } r \in [0, r_1) \tag{21}$$
$$r^2\frac{d^2u}{dr^2} + r\frac{du}{dr} - (1+\gamma^2r^2)u = -\frac{5ab}{2}(1+\nu)r^6, \quad \text{on } r \in [r_1, 1]. \tag{22}$$

We solve (21) and (22) with boundary conditions (15), (16) and continuity in cell deformation and stress at the adhesion boundary $r_1$ using the method of variation of parameters. Thus, we determine the analytic solution as

$$u = \begin{cases} \alpha_I r - \frac{(1+\nu)ab}{14}r^6, & r \in [0, r_1) \\ \alpha_O I_1(\gamma r) + \beta_O K_1(\gamma r) + \frac{5(1+\nu)ab}{2\gamma^6}\left(\gamma^4r^4 + 15\gamma^2r^2 - \frac{45\pi}{2}L_1(\gamma r)\right), & r \in [r_1, 1], \end{cases} \tag{23}$$

in terms of modified Bessel and Struve functions[3]. The coefficients $\alpha_I$, $\alpha_O$, and $\beta_O$ are determined from the boundary conditions to be

$$
\begin{aligned}
\alpha_O = \frac{(1+\nu)}{2\gamma} \cdot a \cdot \Bigg( \frac{5b}{\gamma^5} \Bigg[ \frac{G(\gamma)}{g(\gamma r_1)} \left( \frac{1}{7}\gamma^5 r_1^5 + 3\gamma^3 r_1^3 + 15\gamma r_1 - \frac{45\pi}{2} m(\gamma r_1) \right) \\
- \left( (4+\nu)\gamma^3 + 15(2+\nu)\gamma - \frac{45\pi}{2} M(\gamma) \right) \Bigg] - (1+b) \Bigg) \\
\cdot \left( F(\gamma) - \frac{f(\gamma r_1)}{g(\gamma r_1)} G(\gamma) \right)^{-1}, \\
\beta_O = \frac{(1+\nu)}{2\gamma} \cdot a \cdot \Bigg( \frac{5b}{\gamma^5} \Bigg[ \frac{F(\gamma)}{f(\gamma r_1)} \left( \frac{1}{7}\gamma^5 r_1^5 + 3\gamma^3 r_1^3 + 15\gamma r_1 - \frac{45\pi}{2} m(\gamma r_1) \right) \\
- \left( (4+\nu)\gamma^3 + 15(2+\nu)\gamma - \frac{45\pi}{2} M(\gamma) \right) \Bigg] - (1+b) \Bigg) \\
\cdot \left( \frac{g(\gamma r_1)}{f(\gamma r_1)} F(\gamma) - G(\gamma) \right)^{-1}, \\
\alpha_I = \frac{1}{r_1} \left( \alpha_O I_1(\gamma r_1) + \beta_O K_1(\gamma r_1) \right) \\
+ \frac{5(1+\nu)ab}{2\gamma^5} \left( \frac{1}{35}\gamma^5 r_1^5 + \gamma^3 r_1^3 + 15\gamma r_1 - \frac{45\pi}{2} \cdot \frac{1}{\gamma r_1} L_1(\gamma r_1) \right),
\end{aligned}
$$

and for simplicity we have defined the functions

$$
\begin{aligned}
F(z) &:= I_0(z) + \frac{(\nu-1)}{z} I_1(z), & f(z) &:= I_0(z) - \frac{2}{z} I_1(z), \\
G(z) &:= K_0(z) - \frac{(\nu-1)}{z} K_1(z), & g(z) &:= K_0(z) + \frac{2}{z} K_1(z), \\
M(z) &:= L_0(z) + \frac{(\nu-1)}{z} L_1(z), & m(z) &:= L_0(z) - \frac{2}{z} L_1(z).
\end{aligned}
$$

We substitute (23) into (5) to evaluate the integral expression for active strain energy

$$
\begin{aligned}
W_{CA} &= \frac{1}{2}\int_A \sigma^A_{ij}\varepsilon_{ij}dA \\
&= \frac{(1+\nu)\pi a}{2}\int_0^1 \left(1+br^5\right)\left(\frac{du}{dr}+\frac{u}{r}\right)rdr \\
&= \frac{(1+\nu)\pi a}{2}\int_0^{r_1} \left(1+br^5\right)\left(2\alpha_I r - \frac{(1+\nu)ab}{2}r^6\right)dr \\
&\quad + \frac{(1+\nu)\pi a}{2}\int_{r_1}^1 \left(1+br^5\right)\Bigg(\alpha_O\gamma r I_0(\gamma r) - \beta_O\gamma r K_0(\gamma r) \\
&\qquad\qquad + \frac{5(1+\nu)ab}{2\gamma^6}\left[5\gamma^4r^4+45\gamma^2r^2-\frac{45\pi}{2}\gamma r L_0(\gamma r)\right]\Bigg)dr \\
&= \frac{(1+\nu)\pi a}{2}\Bigg(\alpha_I {r_1}^2 + \frac{1}{7}\left(2\alpha_I b - \frac{(1+\nu)ab}{2}\right){r_1}^7 - \frac{(1+\nu)ab^2{r_1}^{12}}{24} \\
&\qquad + \alpha_O\left(I_1(\gamma)-r_1I_1(\gamma r_1)\right) + \beta_O\left(K_1(\gamma)-r_1K_1(\gamma r_1)\right) \\
&\qquad + \frac{5(1+\nu)ab}{2\gamma^4}\Bigg(\frac{b\gamma^2}{2}(1-{r_1}^{10}) + \frac{45b}{8}(1-{r_1}^8) + \gamma^2(1-{r_1}^5) \\
&\qquad\qquad + 15(1-{r_1}^3) - \frac{45\pi}{2\gamma^2}\left(L_1(\gamma)-r_1L_1(\gamma r_1)\right)\Bigg) \\
&\qquad + \int_{\gamma r_1}^{\gamma}\left(\frac{\alpha_0 b}{\gamma^6}z^6I_0(z) - \frac{\beta_0 b}{\gamma^6}z^6K_0(z) - \frac{225(1+\nu)\pi ab^2}{4\gamma^{12}}z^6L_0(z)\right)dz\Bigg).
\end{aligned}
\tag{24}
$$

The final integral expressions in (24) may be explicitly evaluated using integration by parts as

$$
\begin{aligned}
\int z^6 I_0(z)dz &= -5\left(z^5+15z^3-\frac{45\pi}{2}zL_1(z)\right)I_0(z) \\
&\quad + \left(z^6+25z^4+225z^2-\frac{225\pi}{2}zL_0(z)\right)I_1(z), \\
\int z^6 K_0(z)dz &= -5\left(z^5+15z^3-\frac{45\pi}{2}zL_1(z)\right)K_0(z) \\
&\quad - \left(z^6+25z^4+225z^2-\frac{225\pi}{2}zL_0(z)\right)K_1(z), \\
\int z^6 L_0(z)dz &= \frac{5}{\pi}\left(\frac{1}{3}z^6+\frac{15}{2}z^4+45z^2\right) - 5\left(z^5+15z^3+45z\right)L_0(z) \\
&\quad + \left(z^6+25z^4+225z^2\right)L_1(z) + \frac{225z^2}{\pi}\,{}_2F_3\left(1,1;\frac{3}{2},\frac{3}{2},2;\frac{z^2}{4}\right),
\end{aligned}
$$

where ${}_pF_q(a_1, ..., a_p; b_1, ..., b_q; z)$ is the generalised hypergeometric function [3, 4] used to express $\int zL_1(z)dz$.

The substrate strain energy is evaluated by substituting (23) into expression (6) to obtain

$$
\begin{aligned}
W_S &= \frac{1}{2}\int_{A_{ad}} \gamma^2 \mathbf{u}\cdot\mathbf{u} dA \\
&= \gamma^2 \pi \int_{r_1}^{1} u^2 r dr \\
&= \pi \int_{\gamma r_1}^{\gamma} \left( \alpha_O I_1(z) + \beta_O K_1(z) + \frac{5(1+\nu)ab}{2\gamma^6} \left( z^4 + 15z^2 - \frac{45\pi}{2} L_1(z) \right) \right)^2 z dz \\
&= \pi \Bigg( {\alpha_O}^2 \int_{\gamma r_1}^{\gamma} I_1(z)^2 z dz + {\beta_O}^2 \int_{\gamma r_1}^{\gamma} K_1(z)^2 z dz + 2\alpha_O \beta_O \int_{\gamma r_1}^{\gamma} I_1(z) K_1(z) z dz \\
&\quad + \frac{5(1+\nu)ab}{\gamma^6} \Bigg[ \alpha_O \int_{\gamma r_1}^{\gamma} \left( z^5 + 15z^3 - \frac{45\pi}{2} z L_1(z) \right) I_1(z) dz \\
&\qquad + \beta_O \int_{\gamma r_1}^{\gamma} \left( z^5 + 15z^3 - \frac{45\pi}{2} z L_1(z) \right) K_1(z) dz \Bigg] \\
&\quad + \frac{25(1+\nu)^2 a^2 b^2}{4\gamma^{12}} \int_{\gamma r_1}^{\gamma} \Bigg( z^9 + 30z^7 + 225z^5 + \left(\frac{45\pi}{2}\right)^2 z L_1(z)^2 \\
&\qquad - 45\pi z^5 L_1(z) - 675\pi z^3 L_1(z) \Bigg) dz \Bigg). \qquad (25)
\end{aligned}
$$

We may explicitly evaluate the expression (25) using the integrals

$$
\begin{aligned}
\int zI_1(z)^2dz &= zI_0(z)I_1(z) - \frac{1}{2}z^2\left(I_0(z)^2 - I_1(z)^2\right),\\
\int zK_1(z)^2dz &= -zK_0(z)K_1(z) - \frac{1}{2}z^2\left(K_0(z)^2 - K_1(z)^2\right),\\
\int zI_1(z)K_1(z)dz &= \frac{1}{2}z^2\left(I_0(z)K_0(z) + I_1(z)K_1(z)\right) - zI_1(z)K_0(z),\\
\int z^5I_1(z)dz &= \left(z^5 + 15z^3 - \frac{45\pi}{2}zL_1(z)\right)I_0(z) - \left(5z^4 + 45z^2 - \frac{45\pi}{2}zL_0(z)\right)I_1(z),\\
\int z^3I_1(z)dz &= \left(z^3 - \frac{3\pi}{2}zL_1(z)\right)I_0(z) - 3\left(z^2 - \frac{\pi}{2}zL_0(z)\right)I_1(z),\\
\int zL_1(z)I_1(z)dz &= \frac{1}{\pi}z^2I_1(z) - \frac{1}{2}\left(z^2L_0(z)I_0(z) - z^2L_1(z)I_1(z) - 3zL_1(z)I_0(z) + zL_0(z)I_1(z)\right),\\
\int z^5K_1(z)dz &= -\left(z^5 + 15z^3 - \frac{45\pi}{2}zL_1(z)\right)K_0(z) - \left(5z^4 + 45z^2 - \frac{45\pi}{2}zL_0(z)\right)K_1(z),\\
\int z^3K_1(z)dz &= -\left(z^3 - \frac{3\pi}{2}zL_1(z)\right)K_0(z) - 3\left(z^2 - \frac{\pi}{2}zL_0(z)\right)K_1(z),\\
\int zL_1(z)K_1(z)dz &= \frac{1}{\pi}z^2K_1(z) + \frac{1}{2}\left(z^2L_0(z)K_0(z) + z^2L_1(z)K_1(z) - 3zL_1(z)K_0(z) - zL_0(z)K_1\right.\\
\int \left(z^9 + 30z^7 + 225z^5\right)dz &= \frac{1}{10}z^{10} + \frac{15}{4}z^8 + \frac{75}{2}z^6,\\
\int zL_1(z)^2 &= zL_0(z)L_1(z) + \frac{1}{2}z^2\left(L_1(z)^2 - L_0(z)^2\right)\\
&\quad + \frac{2}{\pi}\left(z^2L_1(z) - 2zL_0(z) + \frac{2z^2}{\pi} + \frac{2z^2}{\pi}\,{}_2F_3\left(1,1;\frac{3}{2},\frac{3}{2},2;\frac{z^2}{4}\right)\right),\\
\int z^5L_1(z)dz &= -\frac{1}{\pi}\left(\frac{1}{3}z^6 + \frac{15}{2}z^4 + 45z^2\right) + \left(z^5 + 15z^3 + 45z\right)L_0(z)\\
&\quad - 5\left(z^4 + 9z^2\right)L_1(z) - \frac{45z^2}{\pi}\,{}_2F_3\left(1,1;\frac{3}{2},\frac{3}{2},2;\frac{z^2}{4}\right),\\
\int z^3L_1(z)dz &= -\frac{1}{2\pi}\left(z^4 + 6z^2\right) + \left(z^3 + 3z\right)L_0(z) - 3z^2L_1(z) - \frac{3z^2}{\pi}\,{}_2F_3\left(1,1;\frac{3}{2},\frac{3}{2},2;\frac{z^2}{4}\right).
\end{aligned}
$$

As in the case of complete adhesion, we can see that the expressions (24) and (25) are proportional to $a^2$ and quadratic in $b$ which may be used to determine the differential contractility profile required to maintain a particular constraint on either $W_{CA}$ or $W_S$.

### S2 Numerical methods for case of cell adhered in distributed patches

When considering adhesions distributed in discrete spots, numerical solutions are required to determine the deformations. The two geometries considered were 20 circular spots distributed evenly or in two polarised clusters around the cell edge. Each spot has radius $r_s$, and radial position $r_p = 0.98r_0 - r_s$. For an even distribution of spots, circles of adhesion were placed at angles $2\pi/20$. For two adhesion clusters at opposite poles of the cell, the angle between adjacent spots in the same cluster was given by $2\arcsin((r_s + 0.015r_0)/r_p)$. Each cluster had 10 spots, and the starting spots of each cluster were separated by angle $\pi$. For each case, a mesh was generated automatically from the defined geometry using the generateMesh command within the MATLAB PDE Toolbox (with `Hmax`, maximum edge length, set at 0.02).

To numerically solve the force balance equation (1) the PDE Toolbox in MATLAB (R2018a) was used. The force balance equation may be expressed as a general elliptic PDE in two dimensions

$$\begin{aligned}
-\nabla \cdot (c_{11}\nabla u_1) - \nabla \cdot (c_{12}\nabla u_2) + a_{11}u_1 + a_{12}u_2 &= f_1 \\
-\nabla \cdot (c_{21}\nabla u_1) - \nabla \cdot (c_{22}\nabla u_2) + a_{21}u_1 + a_{22}u_2 &= f_2.
\end{aligned}$$

As in the radially symmetric cases of a complete adhesion or an adhered ring, we have normalised length scales by the cell radius $r_0$. Thus, specifically the coefficient matrices are

$$c_{11} = \begin{pmatrix} 1 & 0 \\ 0 & \frac{(1-\nu)}{2} \end{pmatrix}, \; c_{12} = \begin{pmatrix} 0 & \nu \\ \frac{(1-\nu)}{2} & 0 \end{pmatrix}, \; c_{21} = \begin{pmatrix} 0 & \frac{(1-\nu)}{2} \\ \nu & 0 \end{pmatrix}, \; c_{22} = \begin{pmatrix} \frac{(1-\nu)}{2} & 0 \\ 0 & 1 \end{pmatrix},$$

$(a_{11}, a_{22}) = T(\mathbf{x})\gamma^2$, and $a_{12} = a_{21} = f_1 = f_2 = 0$ for uniform isotropic contractility. The no stress boundary condition is input as generalised Neumann boundary conditions

$$\mathbf{n}\cdot(c_{11}\nabla u_1)+\mathbf{n}\cdot(c_{12}\nabla u_2)+q_{11}u_1+q_{12}u_2=g_1$$
$$\mathbf{n}\cdot(c_{21}\nabla u_1)+\mathbf{n}\cdot(c_{22}\nabla u_2)+q_{21}u_1+q_{22}u_2=g_2,$$

with $q_{11}=q_{12}=q_{21}=q_{22}=0$ and $(g_1,g_2)=-((1+\nu)P_0/2)\mathbf{n}$. Hence, we see that the problem can be completely parametrised by $\gamma$, $P_0$ and $\nu$ for a general adhesion geometry.

The integrals (5) and (6) for strain energies were calculated from the numerically computed data using the Gaussian quadrature of degree 1 to approximate the integral on each triangle of the mesh. For a general pattern of adhesion, we no longer have an analytic expression for $W_{CA}$ or $W_S$. However, we may show that the cellular deformations, strains and stresses scale linearly with contractile pressure ($P_0$) for a given distribution of cell-substrate adhesions. Hence, $W_{CA}$ and $W_S$ are still proportional to $P_0^2$ as for the radially symmetric adhesion distributions considered previously.

[3] M Abramowitz and I A Stegun. *Handbook of mathematical functions: with formulas, graphs, and mathematical tables*, volume 55. Courier Corporation, 1964.

[4] L J Slater. *Generalized hypergeometric functions.* Cambridge Univ. Press, 1966.
